# Mechanosensory Signaling in Axolotl Courtship and Evolution of Communication

**DOI:** 10.64898/2026.04.13.718269

**Authors:** Taylor M. Rupp, Jeanette M. McGuire, Heather L. Eisthen

## Abstract

In behavioral biology, many models have been proposed to explain how communication systems evolve. Within neuroethology, the principle of sender-receiver matching has spurred much research in auditory communication, but fewer studies on mechanosensory communication. We investigated sender-receiving matching in mechanosensory communication, focusing on the “hula”, a courtship behavior that produces an undulating movement of the tail, in axolotls (*Ambystoma mexicanum*), an aquatic salamander.

We characterized typical courtship behaviors, then quantified tail-motion parameters (speed, sweep angle, and elevation angle) from males as they performed the hula. We then constructed a “Robotail”, a robotic device that mimics the physical and motion properties of the male tail during courtship. Interestingly, females initially responded to the Robotail as if it were a prey item, an effect that was mitigated by the addition of male whole-body odorants. We examined female behavioral responses to changes in individual Robotail movement parameters and found that speed and sweep angle were important to locomotion. Females transitioned between locomotor states more often when exposed to combinations of wide sweep angles and fast speeds from the Robotail, which males perform moderately or rarely, perhaps reflecting a preference for vigorous movements. We then assessed neural responses to stimuli generated by the Robotail by recording from the anterodorsal lateral line nerve (ADLLn), which innervates the mechanosensory neuromasts on the snout. The female ADLLn responded most vigorously when stimulated with moderate sweep angles and speeds, parameters often used by courting males. Thus, our behavioral results support a receiver bias model but our neurophysiological results support a sender-receiver matching model within mechanosensory communication during courtship in axolotls. Our results also provide novel evidence that mechanosensory cues generated during hula behavior in salamanders play a role in courtship.

## Introduction

The evolution of communication systems poses an interesting problem in behavioral biology. How and why do animals detect novel communication signals? And why would an animal produce a signal without assurance that the intended receivers can detect it? Many models have been proposed to address these issues, with some emphasizing the potential role of cues inadvertently produced by senders that provide information to receivers, and others emphasizing the potential role of features of the receiver’s sensory systems in shaping signals produced by senders (Bradbury & Vehrenkamp, 2011). Within the field of neuroethology, the principle of sender-receiver matching, which posits that the features of a signal should complement the features of the detector and vice versa, has attracted much research. This idea, borrowed from engineering, fueled important early work on courtship calling in crickets (Alexander, 1962, Hoy et al., 1977, Mhatre et al., 2011, Ronacher, 2019). In addition to crickets, sender-receiver matching has been demonstrated in some studies of the acoustic properties of birdsong (Gall, 2012; Henry, 2016). In weakly electric fish, sender-receiver matching has been demonstrated to occur in detection of self-generated electrical signals (Zakon & Meyer, 1983) as well as in electrosensory communication (Allen & Marsat, 2019). Sender-receiver matching has been the subject of little investigation in other areas of animal communication.

In many fishes and aquatic amphibians, the lateral line system contains mechanosensory organs called neuromasts. Their role in intraspecific communication is poorly understood. Studies with teleost fishes indicate that some species produce body movements that direct hydrodynamic stimuli towards conspecifics to facilitate schooling (Coombs & Montgomery, 2014) as well as in more direct social interactions, such as aggression (Butler & Maruska, 2015) and mating (Butler & Maruska, 2016). In salmon, the lateral line has been shown to respond to vibratory signals generated by tail movements during courtship (Satou et al., 1991, 1994) but the degree of matching between the frequency and response is unclear.

Here, we investigated sender-receiver matching in axolotls (*Ambystoma mexicanum*), aquatic salamanders that reproduce year-round in the laboratory. The mode of reproduction in most salamanders is unusual among vertebrates: sperm is transferred externally but fertilization occurs internally. Males axolotls deposit spermatophores on the substrate, and the sperm cap that sits on top of the jelly stalk is picked up by the female using her cloaca, a process that requires close coordination of movements in both individuals. During early courtship male axolotls advertise to females by hulaing, a behavior in which the male sways the pelvis side to side, generating undulating tail movements (Verrell, 1982). If the female is receptive the courtship may progress to a stage in which the male begins walking while hulaing with the female following closely behind. Although tail undulations characterize courtship in many salamander species, the communicative function of this behavior has only been empirically demonstrated in *Hynobius leechii* (Kim et al., 2009). Such tail movements in axolotls and other aquatic salamanders have been proposed to function in dispersing chemical cues released from the male’s cloaca (Tinbergen & Pelkwijk, 1938, Salthe, 1967, Verrell, 1984, Houck & Arnold, 2003), but certainly also generate stimuli that could be detected by lateral lines. Further, although the tuning properties of the mechanosensory lateral line in axolotls have been investigated by previous researchers using a small vibrating sphere (Münz et al., 1982), the role of the mechanosensory lateral line in communication remains unexplored.

To evaluate the role of mechanosensory cues in courtship communication and to assess the match between male displays and female preferences, we studied courtship of freely interacting axolotls to quantify aspects of the hula; constructed a robotic tail, a “Robotail”, to simulate hula movements that represent natural and contrived combinations of hula parameters; tested female behavioral responses to the Robotail; and recorded responses in the anterodorsal lateral line nerve (ADLLn) evoked by the Robotail. Behaviorally, females responded to changes in individual hula parameters (sweep angle, speed, and elevation angle) with changes in locomotion and physical contact with the Robotail. In examining responses to combinations of hula parameters, we found that females transitioned between locomotor states most often when exposed to extreme combinations (fast and wide hulas) that males moderately or rarely perform. In contrast, responses in the ADLLn were strongest when stimulated with movement combinations characteristic of natural courtship behavior (moderate hulas). Thus, our neurophysiological results provide support for the hypothesis that sender-receiver matching occurs in mechanosensory signaling in axolotl courtship, broadening the scope of sensory modalities to which this important principle applies. In contrast, our behavioral results support models that have been referred to as sensory bias, receiver bias, or perceptual bias, in which female receivers show preferences for male traits that are extreme or even beyond the ability of males to produce (Ryan, 1998; Ryan and Cummings, 2013).

## Materials and Methods

### Subjects

Sexually mature adult axolotls (*Ambystoma mexicanum*) were obtained from the Ambystoma Genetic Stock Center at the University of Kentucky. Animals were housed in ∼114 L aquaria at temperatures between 18 and 22°C and were separated by sex into groups of 1-3. Aquaria were filled with an artificial freshwater solution, Holtfreter’s solution (Armstrong et al., 1989) at 40-100%, supplemented with Replenish solution (Seachem Laboratories, Madison, GA). Axolotls reproduce year-round (Armstrong & Malacinski, 1989; Eisthen & Krause, 2012) and we carried out experiments year-round. We updated the light timers in our facility monthly to match the natural sunrise and sunset in axolotls’ native habitat, Mexico City. Housing protocols and experiments were conducted with the approval of the Institutional Animal Care and Use Committee of Michigan State University (approval numbers 10/15-154-00, PROTO201800106).

### Documenting behavior from live axolotl courtship

To establish a baseline for quantifying courtship behaviors in axolotls, we recorded 30 trials with interacting pairs of axolotls (n = 10 males, 19 females). Each male was paired with 3 different females with the caveats that no female was used in two trials in a row and females that laid eggs were not used again. For each trial, we placed a male and a female in a ∼114 L aquarium (approximately 90 cm long x 45 cm wide x 30 cm wide) filled with Holtfreter’s solution at the same concentration as their home aquaria and used Sony Nightshot camcorders (model: CMOS) to record top-down and side views. We placed a lightbox equipped with a series of waterproof infrared LED lights underneath the aquarium to illuminate the recording area. We also placed two infrared lamps behind the aquarium and attached a layer of light-colored paper to the outer surfaces of the aquarium to diffuse the light. We believe that the behavior of our study animals was not affected by our use of infrared lighting: measurements of absorbance spectra of isolated rod cells from a closely related species, the tiger salamander (*Ambystoma tigrinum*), indicate an upper limit shorter than IR wavelengths (Cornwall et al., 1984).

Because axolotls are nocturnal, we placed the pair together shortly before the lights in our animal facility turned off for the evening and allowed the pair to interact overnight. We found that courting pairs were most active during the first 2 hrs after the lights turned off and used a conservative time window of 2.5 hrs for analyses of courtship behavior.

Axolotl courtship and mating behaviors are illustrated in **Figure 1**. We used Shoop’s (1960) description of courtship in mole salamanders (*Ambystoma talpoideum*) to guide an initial behavioral survey, then modified some definitions and added others to the ethogram. We organized behaviors into three categories: female locomotion, courtship and mating, and aggression. We quantified the behaviors in our ethogram for all 30 trials and used these definitions, with minor modifications, in trials with the Robotail (**Table 1**).

**Figure 1.**
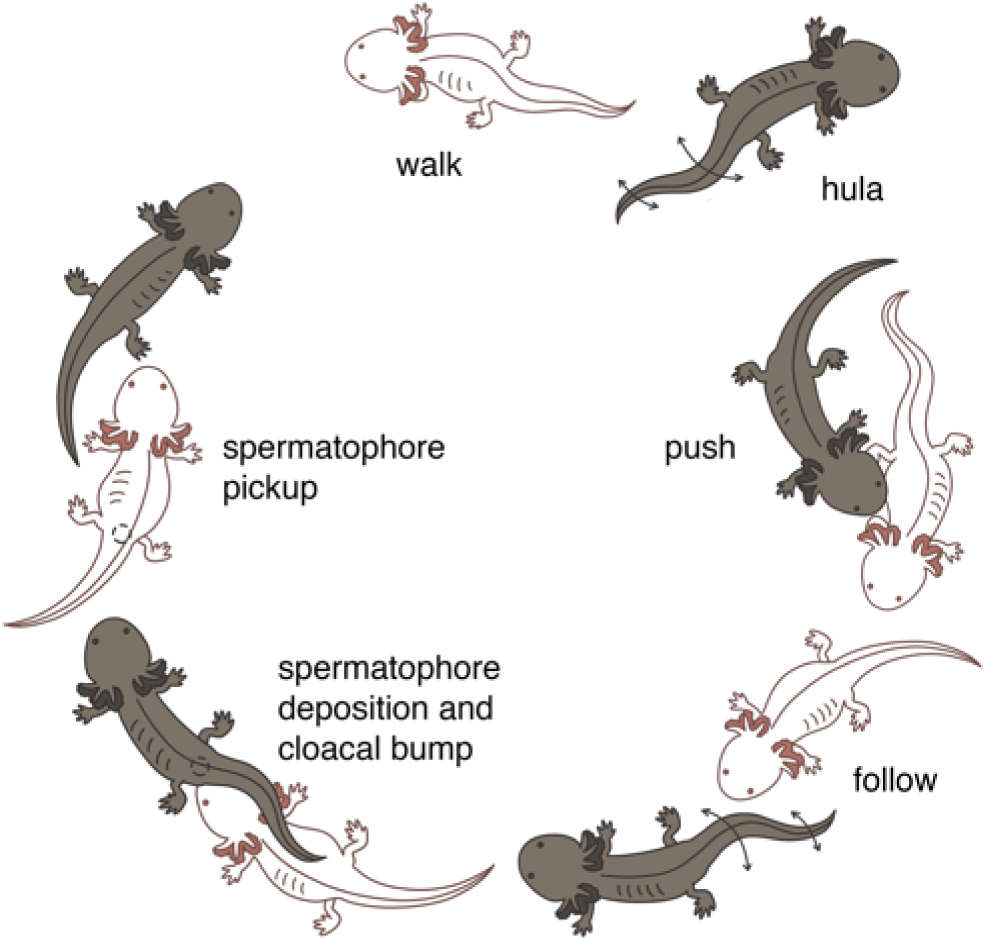
Courtship and mating behavior of male (darker figure) and female (lighter figure) axolotls. Most salamanders use a mating strategy that is unusual among vertebrates in which sperm is transferred externally via a spermatophore, which the female uses to fertilize the eggs internally. Thus, close communication and coordination between male and female are essential for successful mating. Both sexes perform the hula behavior, but courtship often begins with the male hulaing at some distance from the female and frequently approaching to push the female on the flank. Illustration by Ayley Shortridge. The full ethogram of courtship and mating behaviors is presented in **Table 1**.

**Table 1:** Ethogram of male and female courtship behaviors. . We classified behaviors into three categories: female locomotion, courtship and mating, and aggression. Behaviors noted with * were also relevant for trials with the Robotail.

| Behavior |  | Description |
| --- | --- | --- |
| Female<br>*locomotion* | Walk* | Female walks forward or backward at least 3 steps, with all four legs moving. |
|  | Swim* | Female swims forward using her tail, with legs pressed against the body. |
|  | Pause* | Female does not walk or swim. |
|  | Locomote / pause near male* | Female walks, swims, or pauses with head within one male tail length of the male. |
|  | Orient* | Female's head is within one male tail length of the male and she turns to face the male. |
| Courtship and Mating | Push | Male pushes against the female's torso or head with snout. |
|  | Hula* | Animal sways hips and hind legs while undulating tail. |
|  | Male close hula | Male performs hula within one tail length of the female. |
|  | Male distant hula | Male performs hula at least one tail length away from the female. |
|  | Female close hula | Female performs hula within one tail length of the male. |
|  | Female distant hula | Female performs hula at least one male tail length away from the male. |
|  | Follow | Female walks closely behind the male. Starts when female snout is within one head-length of the male's cloacal gland and ends when the male cloacal gland is at least one tail length away from the female's snout. |
|  | Male bump | Male steps backward and touches his cloacal gland or proximal half of the tail to the female's snout. |
|  | Nudge base* | Female steps forward and touches the male's cloacal gland or proximal half of the tail with her snout. |
|  | Nudge tip* | Contact is made between the female's snout and distal half of the male's tail. May be initiated by male or female. |
|  | Deposit | Male deposits a spermatophore onto the substrate. |
|  | Pick up | Female picks up a spermatophore with her cloacal gland. |
| Aggression | Nip* | Female bites the male's tail and immediately releases it. |
|  | Bite* | Female bites the male's tail and briefly keeps jaws engaged. |
|  | Attack* | Female bites the male's tail and thrashes head side to side. |

Because we considered “following” to be a courtship behavior, we only scored female locomotor behaviors when not following a male. We also did not independently quantify male hulaing during following because it consistently co-occurs with following; however, female hulaing that occurred during following was quantified as close hulaing. The females in our study would also cease locomoting for highly variable amounts of time (0.25 sec up to 2.5 hr), and although the term “pause” is typically reserved for short bouts of immobility (Kramer & McLaughlin, 2001) we used “pause” to signify that the female was not locomoting. Finally, we used the terms “near” or “close” to refer to behaviors occurring within 1 tail length of each other and “distant” to refer to events when the pair was more than 1 tail length apart.

We quantified axolotl courtship behaviors using Behavioral Observation Research Interactive Software (BORIS; version 7.9.RC1;Friard & Gamba, 2016). We also recorded the number of spermatophores present in the aquarium and whether the female laid eggs following a trial.

### Quantification of hula motion parameters

We analyzed hula motion parameters for 6 males, 2 that spawned with females during behavioral trials and 4 selected at random. For each male, we only analyzed trials that contained at least one bout of close hulaing with a female that was at least 2 min long and/or a bout of distant hulaing that was at least 30 sec long; thus, we included 3 trials for 1 male, 2 trials for 3 males, and a single trial for the remaining 2 males. For each trial, we analyzed footage during the first 2 hrs after the lights turned off to quantify hula parameters.

We used a modified version of Altmann’s (1974) instantaneous sampling method to analyze male tail movements within each hula bout. For close hula bouts (22 bouts across 4 males), we divided each bout into four 30-sec windows and took measurements for the first 10 sec of each window; for distant bouts (31 bouts across 5 males), which were generally shorter, we analyzed two 15-sec windows and took measurements for the first 5 sec of each one. We used Kinovea software version 0.8.27 (www.kinovea.org) to measure tail sweep angles and frequencies in tandem from the overhead perspective; we watched video footage frame by frame and recorded the lateral angle whenever the proximal third of the tail changed direction. We recorded the maximum and minimum sweep angle for each measurement period and calculated the average minimum and average maximum for each individual. The frequency of tail motion was calculated by dividing the total number of angle measurements by the duration of the measurement period. Elevation angles were measured from the side view by measuring the ventral edge of the male’s tail relative to the aquarium floor. We recorded the maximum and minimum elevation angles during each measurement period and calculated the average value for each measurement.

### Construction of the Robotail, a hula-mimicking robot

To produce hula-like motion in a controlled and repeatable fashion, we constructed a robotic tail, or “Robotail” (**Fig. 2**). The Robotail consisted of a silicone tail, 3D printed components to mount the silicone tail in the aquarium, and electronic components to control the tail’s motions. All components in contact with aqueous solutions were made of non-metallic materials to prevent metal ions from leaching into the Holtfreter’s solution, which can be harmful to salamanders (Bazar et al., 2009), and to prevent the generation of cues that might stimulate the electrosensory lateral line system (Münz et al., 1984).

**Figure 2.**
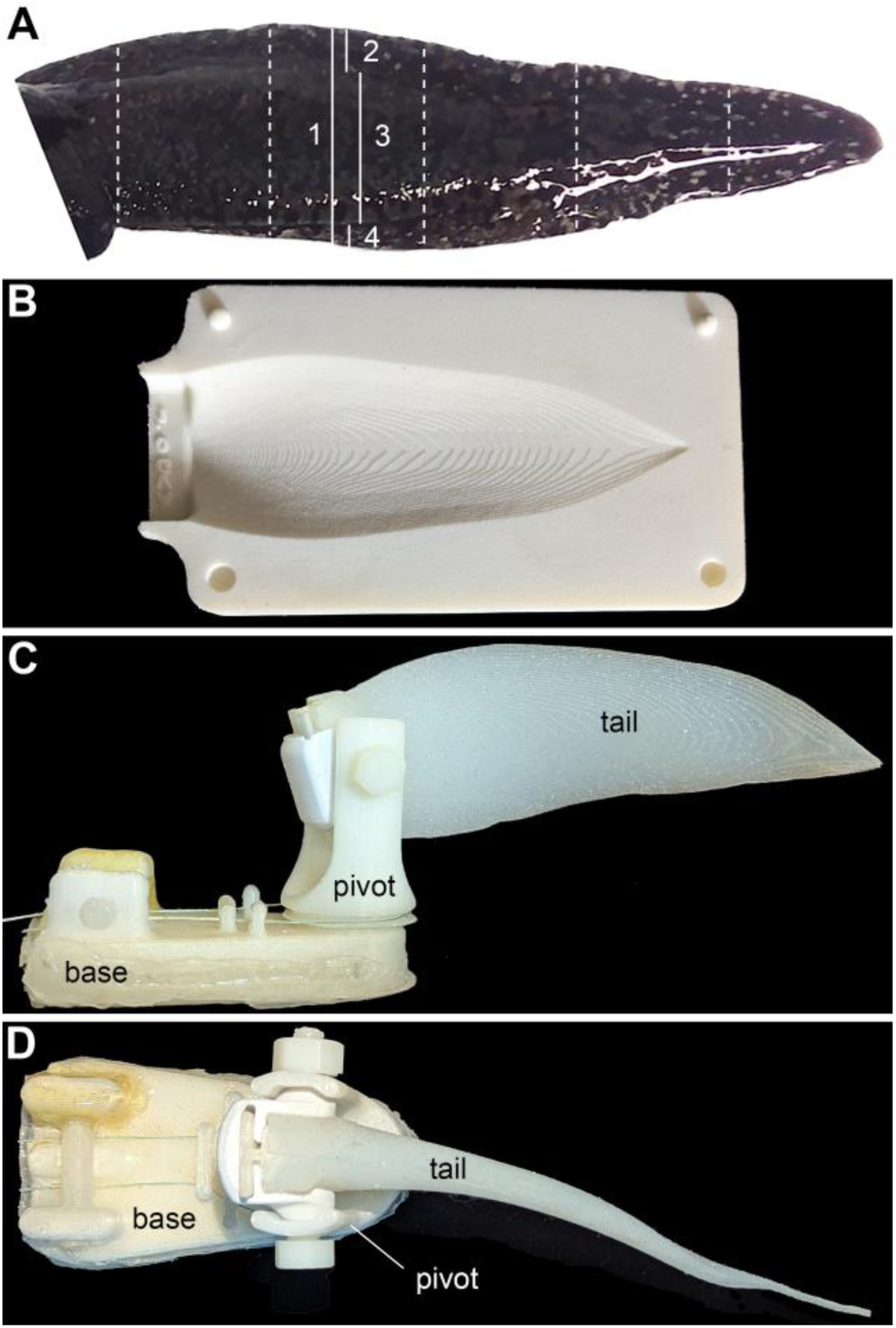
Robotic tail (“Robotail”) used in behavioral and neurophysiological experiments. (**A**) Measurements taken from male axolotls to establish Robotail dimensions. Dashed lines indicate the 5 positions along the tail at which measurements were taken of the (1) total height, (2) dorsal fin width and height, (3) width at the middle of the tail, and (4) ventral fin width and height. (**B**) One side of the mold created to cast the silicone for the tail portion of the Robotail. (**C**, **D**) The silicone tail and mounting components in overhead and side view, respectively. Parts included a silicone tail with embedded plastic clip inserted into a pivoting cylinder mounted on a plastic base. The elevation angle was adjusted using a nylon bolt. Plastic fishing lines controlled side-to-side movement of the pivot.

To obtain measurements necessary to create the silicone tail, we briefly anaesthetized 5 adult male axolotls in 0.1% pH-corrected MS-222 in Holtfreter’s solution; we measured tail parameters from the lightest and heaviest males in our colony and 3 additional randomly-chosen males. The total length of the tail (posterior edge of cloacal gland to tail tip) was divided into 5 equal segments; at each landmark, we measured the width and height of the dorsal and ventral fins, the width at the middle of the tail, and the total height (**Fig. 2A**). We used the median value of each measurement to generate a representative tail, which we modeled, along with a 2-part mold (**Fig. 2B**), in SolidWorks software (version: SolidWorks 2016; Dassault Systèmes, Vélizy-Villacoublay, France). We made the silicone tail from Ecoflex 00-30 (Smooth-On, Inc., Macungie, PA) using a 3D print of our custom mold from Shapeways (Livonia, MI); during the casting process, we embedded a 3D printed clip in the proximal end to attach the tail to the mounting components.

As illustrated in **Figures 2C** and **D**, the mounting components consisted of a 3D printed triangular base plate and a pivoting cylinder with a slot to hold the tail in place. The triangular base was mounted to the aquarium floor using waterproof silicone. We also sourced a silicone salamander toy resembling an ambystomid salamander, removed the tail, and mounted the remainder of the toy in front of the base plate to resemble the shape of a male axolotl. Finally, we attached EVA foam tiles to the aquarium floor to make it flush with the mounting plate and rubber salamander.

We used a series of electronic components to control the oscillating motion of the silicone tail, including a power distribution channel, Victor SP speed controller, 104:1 NeveRest Sport gearbox, NeveRest gearmotor, and Thrifty Throttle 3/Cypress PSoC4™ microcontroller from AndyMark Inc. (Kokomo, IN) as well as a PSoC4™ MiniProg3 programmer (Infineon Technologies AG, Munich, Germany) and a potentiometer (DigiKey, Thief River Falls, MN). A metal gimbal controlled the lateral position of the tail. We wrote custom software in C (Kernighan & Ritchie, 2002) using PSoC™ Creator (Infineon Technologies AG, Munich, Germany) to control the speed and sweep angle of tail oscillation. The elevation angle of the silicone tail was changed manually. Nylon fishing line was used to connect the pivoting cylinder to the gimbal, which was outside of the aquarium. The pivoting motion of the cylinder transferred an undulating motion to the silicone tail, thus simulating the hula of a male axolotl.

### Female behavioral responses to the Robotail

To assess female behavioral responses to both natural and contrived hula motion patterns, we programmed the Robotail to perform 27 hula motion combinations based on the minimum, median, and maximum values for each parameter: thus, we tested sweep angles of 10°, 30°, and 90°, speeds of 0.5 Hz, 1.0 Hz, and 1.5 Hz, and elevation angles of 0°, 20°, and 55°. We tested each motion combination with 6 different females for a total of 162 trials with the Robotail. All trials were conducted after lights out in our animal facility; between trials, we wore headlamps with red lights to minimize disrupting our subjects. All trials were recorded from an overhead angle using a Lorex Night Vision Security Camera system (model: HDIP82W). We filled the test aquarium with Holtfreter’s solution at the same concentration as the home aquarium, placed a female in the aquarium with the Robotail, began each trial with a 5 min acclimation period, and then turned on the Robotail and allowed the female to interact with it for 15 min. We quantified all behaviors defined in **Table 1**.

In preliminary trials we found that female axolotls attacked the Robotail (**Supplemental Video 1**) and reasoned that perhaps the Robotail was perceived as prey rather than a male conspecific. To alleviate this problem, for subsequent trials we introduced male odorants into the aquarium. To do so, we collected whole-body odorants by placing 3 adult males into individual plastic bowls filled with 1 L of Holtfreter’s solution for roughly 24 hr before each trial and combined the contents to minimize effects of individual variation. Odorants were delivered into the aquarium through a silicone tube mounted near the base of the silicone tail; a peristaltic pump delivered odorant solution at a rate of approximately 33 mL/min while the Robotail was on. With the male odorants supplied, bites and attacks were greatly reduced. Between each trial we replaced the Holtfreter’s solution in the aquarium to remove any odorants.

### Electrophysiological experiments

To assess neural responses to stimuli produced by the Robotail, we performed multi-unit recordings from the ADLLn, which innervates the portion of the lateral line sensors on the dorsal snout (Northcutt, 1992), the region that is immediately below the male’s tail when the female is following the male. We tested 3 sweep angles (10°, 30°, 90°) paired with 3 speeds (0.5 Hz, 1.0 Hz, 1.5 Hz) for a total of 9 motion combinations and rotated the order of stimulus presentation across animals. Because we consistently observed that speed and sweep angle affected female behavior but that the effect of elevation angle was minimal, we used a constant elevation angle of ∼45° for our recordings. This angle also minimized interference of the Robotail with our recording electrodes. The testing protocol, which we ran in triplicate for each combination, included a preparation period of 15 sec followed by a stimulus period of 30 sec, with an intertrial interval of 2 min. We analyzed data from recordings with robust action potentials (APs; generally above 1 mV) and stopped recording when responses dropped below 0.5 mV. Using these criteria, our preparations generally remained viable for a period of 3-4 hr and we obtained useable recordings from 14 female axolotls. For comparison, we also obtained recordings from 5 adult males.

Experiments were conducted inside a Faraday cage in a glass aquarium (71 cm x 22 cm x 18 cm) filled to a depth of ∼15 cm with Holtfreter’s solution at the same concentration as the animals’ home aquaria. The mounting plate for the Robotail was attached to the aquarium floor with waterproof silicone and all electronics were housed outside the Faraday cage (**Fig. 4A**).

We rapidly decapitated each subject and exposed the superficial ophthalmic ramus of the right ADLLn by removing a ∼1 cm^2^ piece of skin posterior to the eye. To mimic courtship postures and maximize stimulation of the neuromasts innervated by the ADLLn, the preparation was placed underneath the Robotail such that the anterior edge of the snout was ∼4 cm from the mounting plate and ∼8 cm from the tail tip. A pair of waterproof electrodes with long shafts allowed us to record electrical activity deep under water; an Ag/AgCl wire suction electrode with a borosilicate glass pipette tip was used for recording and an Ag/AgCl pellet electrode acted as the reference electrode. We performed multiunit recordings by lowering the recording electrode onto the surface of the nerve as posteriorly as possible and drawing gentle suction, moving the recording site anteriorly when the signal deteriorated. The reference electrode was submerged in the Holtfreter’s solution approximately 1 cm away from the tip of the recording electrode.

Signals were amplified through a differential amplifier (DP-304, Warner Instruments, Holliston, MA), high-pass filtered (60 Hz), and digitized using a Digidata 1550A digitizer (Molecular Devices, San Jose, CA). We used pCLAMP 10 software (Molecular Devices) with Clampex and Clampfit programs (version 10.7.0.3) for data recording and analysis, respectively. Automated spike data extraction was performed using MATLAB code (MathWorks, Natick, MA, version R2023b), which was modified from a classroom handout created by Wagenaar & Wright (2008) for the Neural Systems and Behavior Course at the Marine Biological Laboratory in Woods Hole, MA.

### Data analyses

To define which combinations of hula parameters were natural (i.e., those that male axolotls perform) and which were contrived (i.e., those that males were either unlikely or unable to perform), we reduced all observations of the three continuous variables into 3 discrete categories that span the range of observed motions, yielding 27 combinations of parameters. Thus, we binned the data for sweep angle (< 30°, ≥ 30° - 60° ≤, ≥ 60°), hula speed (< 0.6 Hz, ≥ 0.6 - 1.1 Hz ≤, ≥ 1.1 Hz), and elevation angle (< 20°,≥ 20° - 55° ≤, ≥ 55°) and further divided the data into “close”, “distant”, or “all” hula bouts. We then labeled each combination as occurring “sometimes” if it occurred less often than the median value, or “often” if it occurred more often than the median value; those that were not observed were labeled “never” (**Supplemental Table 1**). Any bouts in which we were unable to measure all 3 parameters were discarded.

To facilitate comparisons of our behavioral and neurophysiology data (**Fig. 5**), we also examined combinations of sweep angle and hula speed, excluding elevation angle. Fixing elevation angles resulted in the loss of fully contrived conditions (i.e., there was no combination of sweep and speed that was “never” performed once elevation differences were removed).

We then ranked each combination by the number of occurrences and divided them into three categories representing combinations that occurred most “often” (n = 33; i.e., the top one third of the observations for each parameter in each category), or “moderately” (n = 34; i.e., middle third of the observations) and “rarely” (n = 33; i.e., the bottom third of the observations).

We calculated summary statistics (mean, standard deviation, minimum, and maximum) using the “psych” package (version 2.4.3; Revelle, 2009) in R (R Core Team, 2020). For continuous variables, the average duration of each bout of activity was calculated with zeros removed (i.e., total duration divided by count). For discrete variables (i.e., nudge base/tip, nip, bite, attack count), zeros were retained in the dataset. Because females were repeated in the dataset an unequal number of times, all female behaviors and all sex-based differences in behaviors were analyzed using general linear mixed-effects models (GLMM) in R using packages “lme4” (version 1.1.35.3; Bates et al., 2015) and “lmerTest” (version 3.1.3; Kuznetsova et al., 2017) with female ID as the random effect and restricted maximum likelihood (REML) set to “true”.

In analyzing female behavioral responses to the Robotail, we examined the three hula motion parameters separately and in combination. Female ID was included as a random effect in all models. Fixed effects (sweep angle, speed, and elevation angle) were evaluated alone and in combination with the restricted maximum likelihood (REML) set to “false” to allow us to compare among models with different fixed effects (Bolker, 2015). Because elevation was not included as a variable in our neurophysiology experiments, we excluded elevation in mixed-model analysis of responses to combinations of parameters. In addition, model comparisons were evaluated using corrected Akaike information criterion (AICc) with delta < 3 considered equally weighted and we relied on rules of parsimony to define the best model. Because the results of the AICc analyses were largely consistent with the results of our mixed model analyses, we do not discuss them further; the results are presented in **Supplemental Tables 2-4**.

We also analyzed female neural responses to the Robotail in relation to hula motion parameters both separately and in combination. As our dependent variable, we measured the number of action potentials per second (firing rate or FR) from ADLLn recordings. To standardize among individuals and to account for higher FRs observed in males compared to females, we calculated the percent change in FR for each trial as ((Stimulation FR – Preparation FR)/Preparation FR)*100. For example, a trial with a Preparation FR of 5.93 APs/sec and a Stimulation FR of 10.2 APs/sec would constitute a ∼72% percent change. We evaluated percent change in FR with GLMM using the R packages “lme4” (version 1.1.35.3; Bates et al., 2015) and “lmerTest” (version 3.1.3; Kuznetsova et al., 2017) with sweep angle and speed included as fixed effects. To evaluate differences in the percent change in FR between sexes, animal ID was included as a random effect. In sex-specific models male or female ID (as appropriate) was included as a random effect.

## Results

### Live courtship behavior

#### Female locomotion

Summary statistics are presented as mean(±SD) in the text that follows. In each trial, females spent the most time pausing (90(±27) min) followed by locomoting (50(±20) min). Individual bouts of pausing lasted 3(±13.6) min, whereas individual bouts of locomoting lasted 0.2(± 0.1) min. While locomoting, females spent much more time walking (43(±21) min) than swimming (7(±10) min). Females that spent time close to a male spent a total of 24(±10) min, with visits averaging 0.5(±1.4) min in duration. Females oriented towards the male an average of 6(±7) times during a trial. Remaining stationary thus appears to be an important component of the female courtship ritual.

Female locomotion was affected by the presence or absence, but not proximity, of a hulaing male. Females exhibited shorter bouts of walking when males were hulaing compared to when they were not (**Fig. 3A**, GLMM, t = -6.8, p < 0.01). Females also transitioned among walking, swimming, and pausing significantly more often when a male was hulaing versus when there was no hulaing occurring (**Fig. 3D**, GLMM, t = -3.47, p = 0.001); however, the proximity of a hulaing male did not affect females’ transitions between locomotor states (GLMM, t = 0.03, p = 0.98). Thus, the presence of a hulaing male caused females to increase their transitions between locomotor states, but females exhibited similar activity levels whether the male was hulaing close to them or at a distance.

**Figure 3.**
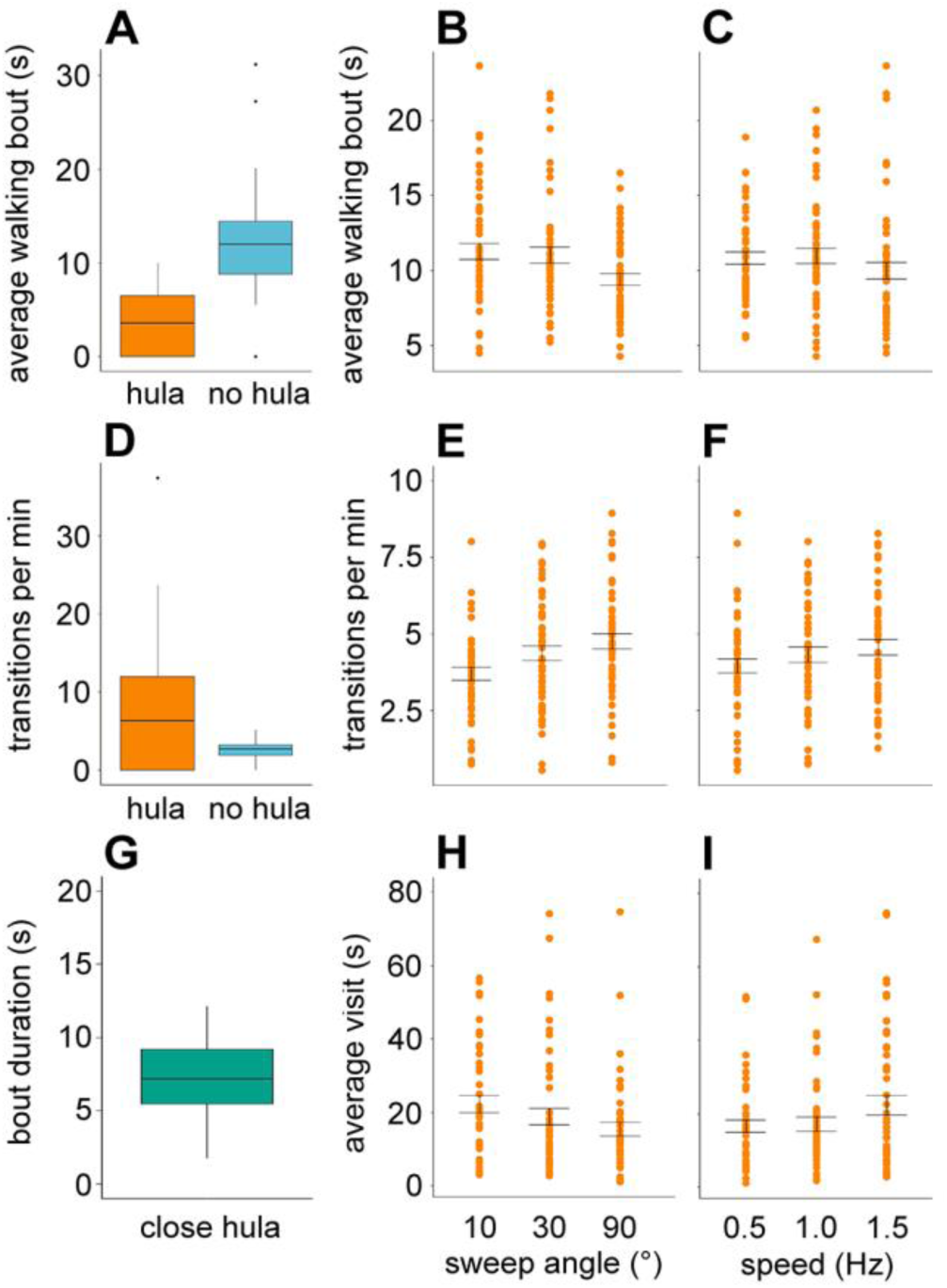
Behavioral responses of female axolotls to live males (left) and to the Robotail (center and right). (**A**) Females exhibited shorter bouts of walking when males were hulaing. Females also engaged in shorter bouts of walking in response to (**B**) the widest sweep angle of the Robotail, and (**C**) the fastest Robotail speed. (**D**) Females transitioned among locomotor states more often when the male was hulaing, (**E**) when presented with sweep angles greater than 10°, and (**F**) when exposed to the fastest Robotail speed. (**G**) Males perform hulas both when a female is close (<1 tail length) or distant (>1 tail length); a close hula is akin to a “visit” of a female to the Robotail. Female visits to the Robotail were of longer duration than with a hulaing male, but visit duration (**H**) decreased when stimulated by the widest sweep angle and (**I**) increased when stimulated by the fastest speed. Panels **A**, **D**, **G**: Each box and whisker plot illustrates the minimum, lower quartile, median, upper quartile, and maximum value; dots represent outlier values and zeroes have been removed. Panels **B**, **C**, **E**, **F**, **H**, **I**: Each dot represents the data from a single individual, and bars represent the standard error of each group. Supporting mixed-model analyses are presented in **Table 2**.

#### Courtship and mating behaviors

Among the 27 pairs of axolotls that courted in our study, males and females spent about half (75±44) min of each 150-min trial engaged in courtship and mating behaviors, suggesting a high level of effort and coordination between the sexes. We found key differences between males and females in both bumping and hula behaviors. Males bumped females (90(±145) significantly more often than females bumped males (51(±71); GLMM, t = 2.45, p = 0.02).

Males and females did not differ substantially in overall amount of time spent hulaing; among individuals that hulaed (10 males, 14 females), males hulaed in total for 8(±8) min and females hulaed for 5(±13) min (GLMM, t = 1.43, p = 0.16). However, the duration of each male hula bout (8.77(±4.5) sec) was longer than that for females (6.04(±2.71) sec; GLMM, t = 2.38, p = 0.02). When in close proximity, males hulaed for more total time than females (males, 5(±4) min, females, 2(±5) min; GLMM, t = 2.45, p = 0.02); however, the duration of each bout of close hulaing was similar for males and females (males, 7.36(±3.03) sec, females, 6.55(±2.33) sec; GLMM, t = 0.87, p = 0.39). At distant ranges, males and females hulaed for a similar amount of time overall (males, 4(±4) min, females, 3(±9) min; GLMM, t = 0.55, p = 0.59); however, males exhibited longer distant hula bouts than females (males, 11.11(±6.05) sec, females, 5.33(±3.07) sec; GLMM, t = 3.80, p = 0.0008).

Males deposited up to 5 spermatophores in 12 out of 30 trials. Due to the video recording angle we were unable to observe females picking up a spermatophore; however, 2 females laid eggs after the trial ended.

#### Aggression

Female aggression towards the male during courtship was minimal and restricted to nipping behavior, which occurred in 27% of our trials and was limited to an average of 1.25 nips per trial.

### Hula motion parameters

During close hulaing, male axolotls generally used moderate sweep and elevation angles and speeds, performing two combinations often: 30° sweep and 20° elevation, at either 1.0 or 1.5 Hz (**Supplemental Table 1**). Close hula bouts that fell into the “sometimes” category generally featured moderate to wide (30° - 90°) sweep angles, moderate speeds (1.0 Hz), and moderate to high elevation angles (20° - 55°). While hulaing at a distance, males performed the following 4 motion combinations often: 10° sweep/1.5 Hz/0° elevation, 30° sweep/1.0 Hz/0° elevation, 30° sweep/1.5 Hz/0°elevation, and 30° sweep/1.0 Hz/20° elevation. Distant hula bouts that fell into the “sometimes” category generally featured narrow to moderate (10° - 30°) sweep angles, moderate speeds (1.0 Hz), and low to moderate (0°- 20°) elevation angles. Thus, the tail motion patterns of close hula bouts were typically wider and higher than distant hula bouts, but males tended to hula with moderate speeds regardless of their proximity to a female.

When we pooled all hula bouts and examined the combinations that occurred, we found that males performed all sweep angles, but generally moderate speeds (1.0 Hz) and low to moderate (0°- 20°) elevation angles. The three motion combinations that males performed often overall were 30° sweep/1.0 Hz/0° or 20° elevation and 30° sweep/1.5 Hz/20° elevation. Six of the combinations we tested in our experiment were never performed by males: 1.5 Hz/55° elevation at any sweep angle; 0.5 Hz/55° elevation at extreme sweep angles (10° or 90°); and 90° sweep/0.5 Hz/0° elevation. Overall, males did not lift their tails high if they were moving at the extremes of their speed range (0.5 or 1.5 Hz) or lower their tails if they were moving slowly and widely.

### Female behavioral responses to the Robotail

#### Physical contacts with the Robotail

Attacking and nipping the Robotail was reduced, but not eliminated, with the inclusion of male whole-body odorants. In 11 of 18 females, we observed 48 nips occurring in 21 of 162 trials; two attacks co-occurred with instances of nipping. In models with a single fixed effect, we found that females nipped at the tail significantly less often as the sweep angle of the Robotail increased (**Table 2**, GLMM, 10° vs 30°; t = -3.00, p = 0.003; 10° vs 90°; t = -2.55, p = 0.01) and marginally less often when the tail was elevated to 20° (**Table 2**, GLMM, t = -1.73, p = 0.09).

**Table 2.** Summary of Mixed Model analyses of changes in female locomotion and behavioral interactions with the Robotail in response to the 3 levels of each hula motion parameter (sweep, speed and elevation), assessed independently. Female ID was included as a random effect in all analyses. Items in bold are statistically significant (p ≤ 0.05) and those in italics are marginally significant (p ≤ 0.10).

| Independent Variable |  | Locomotion (Ave) |  | Pause (Ave) |  | Walk (Ave) |  | Swim (Ave) |  | Average Visit Duration |  |
| --- | --- | --- | --- | --- | --- | --- | --- | --- | --- | --- | --- |
|  |  | t-val | p-val | t-val | p-val | t-val | p-val | t-val | p-val | t-val | p-val |
| Sweep | 30° | -1.58 | 0.12 | -0.70 | 0.49 | -0.20 | 0.84 | -2.21 | **0.03** | -1.04 | 0.30 |
|  | 90° | -2.15 | **0.03** | -1.39 | 0.17 | -2.91 | **<0.01** | -2.21 | **0.03** | -2.2 | **0.03** |
| Speed (Hz) | 1.0 | 0.78 | 0.44 | -1.47 | 0.15 | -0.23 | 0.82 | 0.62 | 0.54 | 0.43 | 0.67 |
|  | 1.5 | -0.97 | 0.34 | -1.79 | *0.08* | -2.27 | **0.03** | -0.78 | 0.44 | 1.90 | *0.06* |
| Elevation | 20° | 0.62 | 0.54 | 0.76 | 0.45 | 1.34 | 0.18 | 0.31 | 0.76 | -0.29 | 0.77 |
|  | 55° | 0.88 | 0.38 | 0.57 | 0.57 | 0.97 | 0.34 | 0.63 | 0.53 | -0.09 | 0.93 |
| Independent Variable |  | Locomotion (Total) |  | Pause Duration (Total) |  | Walk (Total) |  | Swim (Total) |  | Total Visit Duration |  |
|  |  | t-val | p-val | t-val | p-val | t-val | p-val | t-val | p-val | t-val | p-val |
| Sweep | 30° | 0.58 | 0.56 | -0.57 | 0.57 | 1.64 | *0.10* | -0.84 | 0.40 | -0.51 | 0.61 |
|  | 90° | 0.64 | 0.53 | -0.61 | 0.54 | 0.81 | 0.42 | 0.03 | 0.98 | -1.38 | 0.17 |
| Speed (Hz) | 1.0 | 1.22 | 0.22 | -1.14 | 0.26 | 0.72 | 0.48 | 0.62 | 0.54 | 1.48 | 0.14 |
|  | 1.5 | -0.34 | 0.74 | 0.38 | 0.71 | 0.18 | 0.86 | -0.48 | 0.63 | 2.63 | **<0.001** |
| Elevation | 20° | 0.61 | 0.54 | -0.52 | 0.60 | 0.47 | 0.64 | 0.42 | 0.68 | -1.08 | 0.28 |
|  | 55° | -0.04 | 0.97 | 0.09 | 0.93 | -1.02 | 0.31 | 0.80 | 0.43 | -1.30 | 0.20 |
| Independent Variable |  | Transitions per min |  | Orient Count |  | Nudge Tip Count |  | Nudge Base Count |  | Nip Count |  |
|  |  | t-val | p-val | t-val | p-val | t-val | p-val | t-val | p-val | t-val | p-val |
| Sweep | 30° | 2.34 | **0.02** | -1.14 | 0.26 | 0.50 | 0.61 | -1.89 | *0.06* | -3.00 | **<0.01** |
|  | 90° | 4.11 | **<0.001** | -1.14 | 0.26 | 0.25 | 0.80 | -1.49 | 0.14 | -2.55 | **0.01** |
| Speed (Hz) | 1.0 | 1.36 | 0.18 | 0.00 | 1.00 | 0.58 | 0.57 | -0.48 | 0.63 | 0.25 | 0.80 |
|  | 1.5 | 2.23 | **0.03** | 0.38 | 0.71 | 2.11 | **0.04** | 0.45 | 0.65 | 0.15 | 0.88 |
| Elevation | 20° | -0.72 | 0.47 | -1.92 | *0.06* | -1.43 | 0.15 | -0.59 | 0.56 | -1.73 | *0.09* |
|  | 55° | -1.76 | *0.08* | -1.92 | *0.06* | -2.85 | **<0.01** | -0.54 | 0.59 | 0.58 | 0.56 |

Courtship-related interactions were influenced by all three hula parameters when assessed independently. The fastest speed (1.5Hz) elicited more nudging at the tip relative to the slowest speed (**Table 2**, GLMM, t = 2.11, p = 0.04) and the highest elevation (55°) resulted in fewer tip nudges relative to the lowest elevation (**Table 2**, GLMM, t = -2.85, p = 0.005). Females nudged the base of the tail marginally more often when the sweep angle decreased from 30° to 10° (**Table 2**, GLMM, t = -1.89, p = 0.06).

#### Time spent near the Robotail

Results of statistical mixed-model (GLMM) tests of the effects of the each Robotail movement parameter on behavioral responses are presented in **Table 2**. When the Robotail was moving at its fastest speed of 1.5 Hz, females spent significantly more time close to it (GLMM, t = 2.63, p = 0.009) and each visit was for a moderately longer duration (**Fig. 3I**; GLMM, t = 1.9, p = 0.06). In contrast, females spent less time per visit to the Robotail when it was moving at its widest angle of 90° (**Fig. 3H**; GLMM, t = -2.20, p = 0.03). Additionally, we found a marginally significant effect in which females oriented toward the tail fewer times when the Robotail was at a moderate (20°) or higher (55°) angle (**Table 2**; GLMM, 0° vs 20°; t = -1.92, p = 0.06; 0° vs 55°; t = -1.92, p = 0.06).

#### Locomotor behaviors

The individual Robotail parameters we tested did not affect the total duration of time that females spent locomoting (**Table 2**; |t| < 1.22 and p > 0.22 for all parameters) or pausing (|t| < 1.14, p > 0.26) but did affect the duration of each bout of locomoting or pausing. Females walked for significantly shorter bouts of time as the sweep angle increased, with a significant difference between 10° and 90° (**Fig. 3B**; GLMM, t = -2.91, p = 0.004). Females also walked for shorter bouts of time when the tail was moving at the fastest speed compared to the slowest speed (**Fig. 3C**; GLMM, t = -2.27, p = 0.03) and paused for moderately shorter periods when presented with the highest speed (**Table 2**; GLMM, t = -1.79, p = 0.08).

Females transitioned among locomotor states significantly more often when presented with sweep angles higher than 10° (**Fig. 3E**; GLMM 30°, t = 2.34, p = 0.02; 90°, t = 4.11, p < 0.01) or the highest speed (**Fig. 3F**; GLMM, 1.5Hz, t = 2.23, p = 0.03), and transitioned slightly more often when presented with an elevation angle of 55° (GLMM, t = -1.76, p = 0.08).

Females exhibited shorter bouts of swimming as the sweep angle increased (**Table 2**, GLMM, 10° vs 30°; t = -2.21, p = 0.03; 10° vs 90°; t = -2.21, p = 0.03) and shorter bouts of locomotion (walking plus swimming) when stimulated by the widest sweep angle (**Table 2**, GLMM, t = -2.15, p = 0.03). Thus, when analyzed separately, our data revealed that wider sweep angles and faster speeds generally caused females to switch between locomotion and pausing more rapidly than narrower angles and slower speeds.

In addition to examining the effects of changes in single motion parameters, we evaluated the impact of the combination of sweep angle and speed on female locomotor behaviors. Because transitions among locomotor states encompassed multiple types of locomotion and the rate of transitions was influenced by both speed and sweep, we focused on this metric in our combination analyses. Reducing combinations to just speed and sweep angle yielded similar results to the full, three parameter model, but changed the frequency of some behaviors.

Interestingly, one combination, 30° sweep and 1.5 Hz speed, which occurs “moderately” when elevation is not considered, encompasses elevation angles that males either often or never perform at close proximity. Males often performed hulas with 30° sweep and 1.5 Hz speed at 20° elevation, but never performed 30° sweep/1.5 Hz with 0° or 55° elevation. Therefore, in this case of close hulas, the “moderately” category that emerges when the elevation parameter is collapsed represents inclusion of two extreme elevation angles (0° and 55°).

Females transitioned among locomotor states significantly more frequently when presented with a combination of 30°/1.5 Hz (GLMM, t = 2.37, p = 0.02) or 90°/1.0 Hz (GLMM, t = 2.19, p = 0.03), which males moderately perform, and 90°/1.5 Hz (**Fig. 5**; GLMM, t = 4.01, p <0.001), which males rarely perform. In contrast, the number of transitions in response to the combination that males often perform (30°/1.0 Hz) was only marginally significant (GLMM, t = 1.79, p = 0.08). A combination that males rarely perform, 90°/0.5 Hz, was also marginally significant (GLMM, t = 1.69, p = 0.09). Overall, female axolotls transitioned among states more often when the Robotail performed motion combinations featuring moderate to wide sweep angles and moderate to fast speeds.

### Neurophysiological responses of the ADLLn

#### Female vs. male responses

GLMM analyses of neurophysiological responses to the Robotail are summarized in **Table 3**. Overall, we recorded marginally larger responses to stimulation from the Robotail in females compared to males (**Fig. 4 B, C**; GLMM, t = -2.10, p = 0.06). Additionally, we found that females exhibited marginally stronger responses than males did to a stimulus of 1.5 Hz (GLMM, t = - 1.91, p = 0.07). Finally, we found that females exhibited marginally stronger responses than males did to sweep angles of 30° (GLMM, t = -1.81, p = 0.11) and 90° (GLMM, t = -1.55, p = 0.15).

**Table 3.**
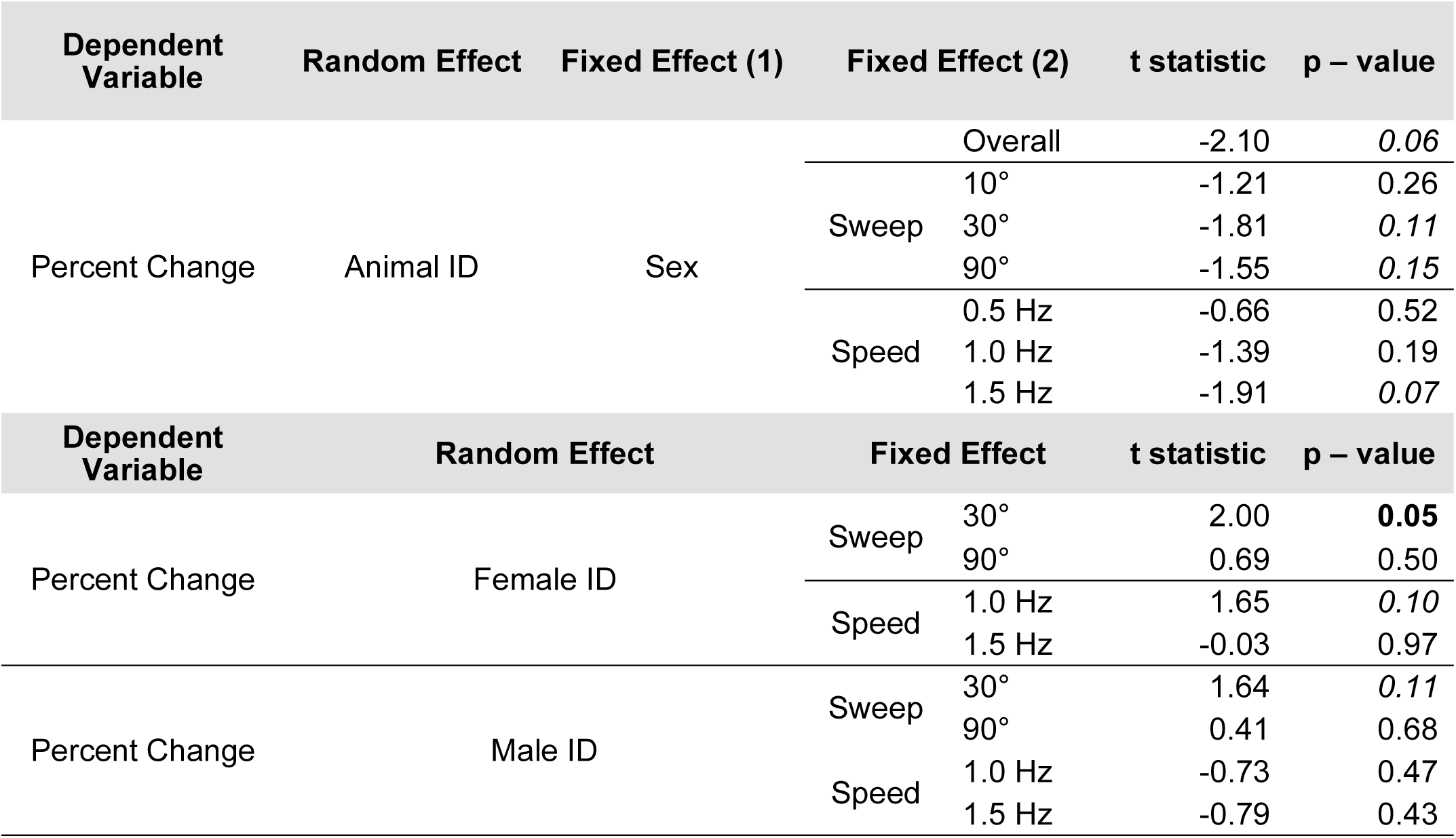
Summary of Mixed Model analyses of percent change in female and male electrophysiological responses to the Robotail stimuli modeling sweep angles and speed. Items in bold are statistically significant (p ≤ 0.05). Those in italics are marginally significant (p ≤ 0.15) but may still be biologically meaningful due to low power from small sample sizes.

**Figure 4.**
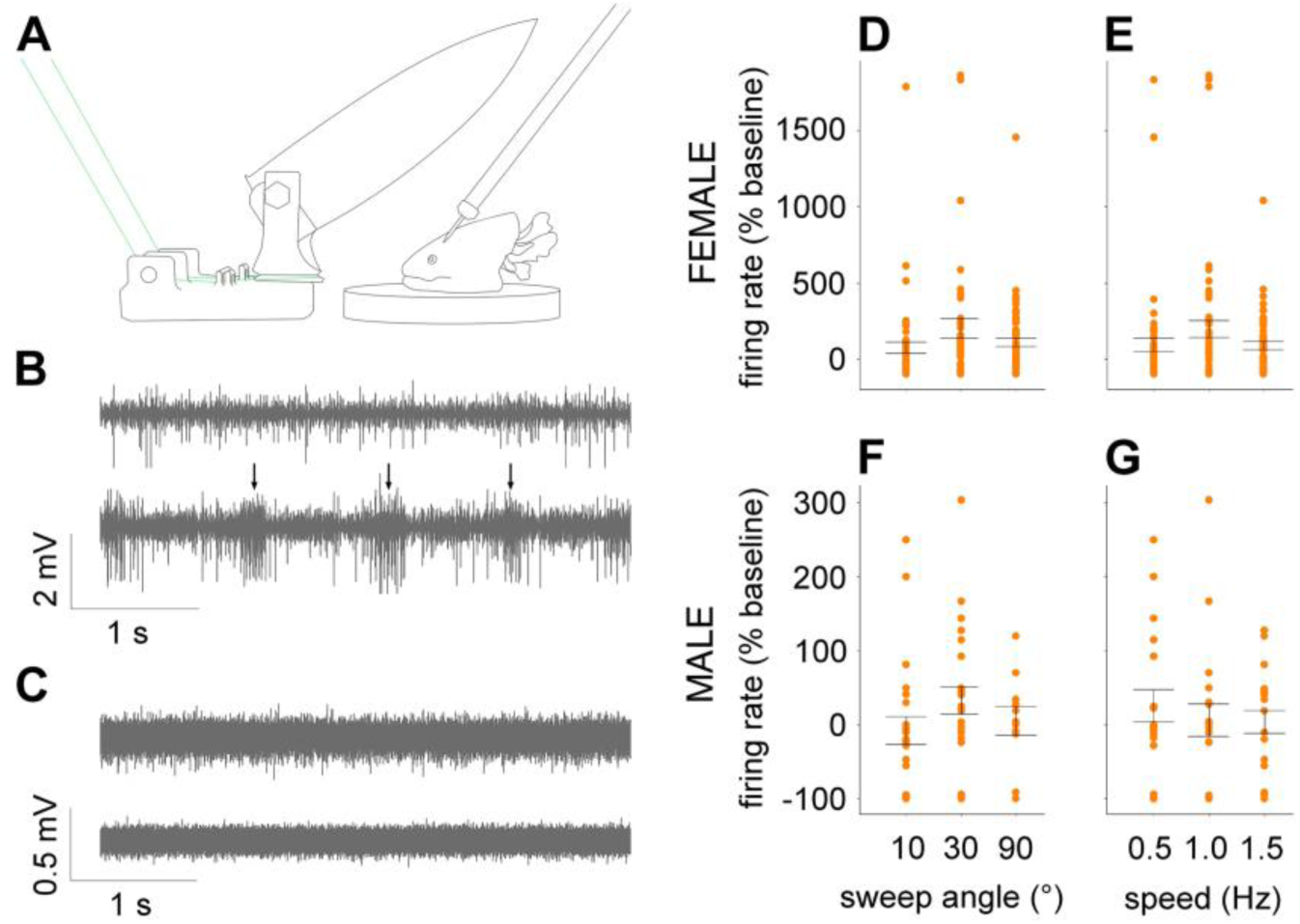
Electrophysiological responses of the anterodorsal lateral line nerve (ADLLn) to stimulation by the Robotail. (**A**) Diagram of the recording setup. The preparation was placed underneath the Robotail to mimic the position of the female relative to the male during mating. The recording electrode was angled such that the tail did not interfere with the electrode while the robot was in operation. (**B**) Representative multi-unit recording from the ADLLn of a female axolotl. Upper trace: baseline firing rate (FR). Lower trace: increased FR in response to stimulation with a 30° sweep angle at 1.0 Hz, with bursts of large action potentials at 1.0 Hz (arrows). (**C**) Representative multi-unit recording from the ADLLn of a male axolotl to the same stimulus. Upper trace, baseline; lower trace, decreased FR in response to the stimulus. (**D**) Change in FR relative to baseline in females in response to stimulation with different sweep angles. A moderate sweep angle of 30° evoked significantly larger excitatory responses than a narrow angle of 10°. (**E**) Firing rate in females did not change in response to stimulation with different speeds. (**F, G**) Change in firing rate relative to baseline in males in response to stimulation with different sweep angles and different speeds, respectively. No significant differences in firing rate were observed. Note the difference in scale of the y-axis in **D** and **E** compared with **F** and **G**. In these panels each dot represents the percent change in FR between preparation and stimulus periods for a given trial and bars represent the standard error for each group. See **Table 3** for supporting mixed-model analyses.

#### Female responses

As illustrated in **Figure 4D**, all 3 sweep angles evoked excitatory responses in the female ADLLn. Independent of speed, a 10° sweep angle increased the firing rate (FR) from 4.33(±4.57) APs/sec during the preparation period to 5.35(±5.88) APs/sec during the stimulation period, an increase of 75.42(±272.48)%. A 30° sweep increased FR 203.75(±461.37)%, and a 90° sweep increased FR 111.05(±223.11)%. We observed a significantly greater increase in FR when the Robotail was moving at a moderate sweep angle of 30° compared to 10° (GLMM, t = 2.00, p = 0.049), but no difference when we presented our preparation with extreme (90°) sweep angles (GLMM, 10° vs 90°, t = 0.69, p = 0.49).

We also recorded excitatory responses to all three speeds (**Fig. 4E**). Independent of sweep angle, we observed the largest increase in FR in response to a 1.0 Hz stimulus (196.24±423.06%); the increase was 94.05(±317.70)% in response to a 0.5 Hz stimulus and 90.80(±196.39)% in response to a 1.5 Hz stimulus. Although responses to the three speeds did not differ significantly, the increase in response was slightly larger with a 1.0 Hz stimulus compared to 0.5 Hz (**Fig. 4E**, GLMM, t = 1.65, p = 0.10).

We also examined responses as a function of combinations of sweep angle and speed. A moderate combination of 30°/1.0 Hz evoked a stronger response than did a narrow, slow combination of 10°/0.5 Hz (**Fig. 5**; GLMM, t = 2.96, p = 0.004), but the same speed (1.0 Hz) with 10° angle elicited a marginally significant result (**Fig. 5**; t = 1.47, p = 0.14). Thus, it appears that moderate combinations evoke the largest responses, and that sweep angle plays an important role determining the degree of excitation in the female ADLLn.

**Figure 5.**
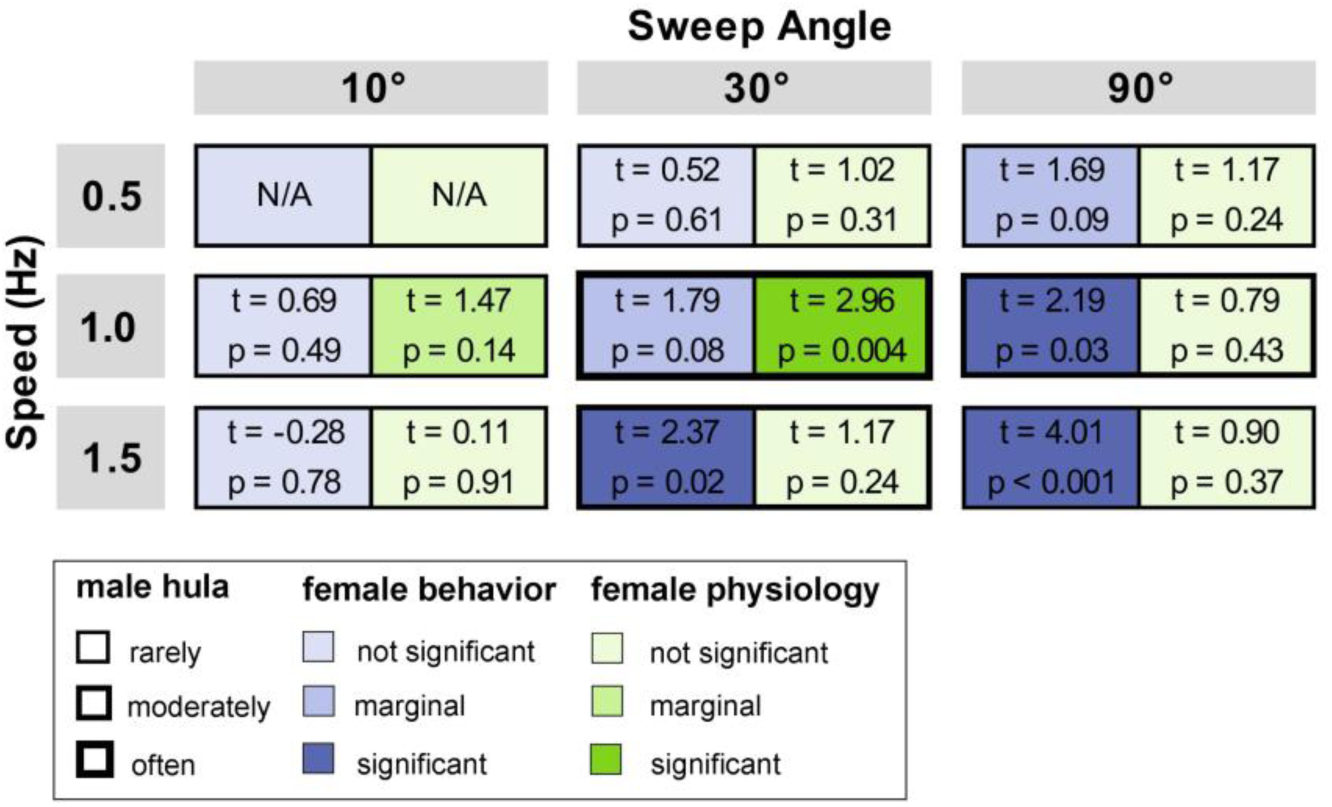
Alignment between male hula behaviors, female behavioral responses to the hula behavior, and responses of the female ADLLn to hula parameters. Legend is shown beneath main experimental results for reference. We found high alignment between male behaviors and female physiological responses, particularly within the 30°/1.0 Hz combination; males performed this combination often and the female ADLLn exhibited a significantly higher FR compared to 10°/0.5 Hz. In contrast, females exhibited a broader behavioral response to the hula combinations we tested, such that females transitioned between locomotor states more often when exposed to wider sweep angles and faster speeds.

#### Male responses

Interestingly, ADLLn activity in males was inhibited when stimulated with a sweep angle of 10°, but excited by wider angles: a 10° stimulus resulted in a decrease in FR of -7.07(±90.14)% but a 30° stimulus caused an increase of 33.00(±91.78)%, and a 90° stimulus resulted in an increase of 5.71(±63.05)%. Although differences in sweep angles independent of speed did not evoke statistically significant percent change in FR, we observed a slightly elevated response to 30° compared to 10° (**Fig. 4F**; GLMM, t = 1.64, p = 0.11). Similarly, we observed inhibitory responses when the Robotail was moving at a slow speed of 0.5 Hz, but excitatory responses to faster speeds, independent of sweep angle: a 0.5 Hz stimulus caused FR to decrease 26.41(±95.16)% but a stimulus of 1.0 Hz resulted in an increase of 6.48(±97.46)% and a 1.5 Hz resulted in an increase of 4.69(±69.35)%. The speed of the Robotail did not affect FR (**Fig. 4G**). Because of our small sample size, we were unable to analyze responses to combinations of parameters.

### Comparison of the male hula to female behavioral and physiological responses

As illustrated in **Figure 5**, we found alignment between male hula behaviors and responses of the female ADLLn to the Robotail. During courtship, male axolotls often hulaed using a combination of 30° sweep angle/1.0 Hz and hulaed a moderate amount with more extreme combinations: 30°/1.5 Hz or 90°/1.0 Hz. When we stimulated the female ADLLn with the Robotail, we found that the nerve exhibited a significantly stronger response to a combination that males often perform, 30°/1.0 Hz, compared to the rarely performed combination 10°/0.5 Hz. As noted above, we also observed a marginally significant excitation of the female ADLLn in response to 10°/1.0 Hz, which males rarely perform.

In contrast, we observed only a moderate level of alignment between male hula patterns and female behavioral responses to the Robotail. Female axolotls transitioned between locomotor states significantly more frequently when the Robotail was oscillating with combinations featuring extreme sweep angles and/or speeds, which males performed either moderately (30°/1.5 Hz; 90°/1.0 Hz) or rarely (90°/1.5 Hz) compared to 10°/0.5 Hz. Females exhibited a marginal increase in transitions/min when stimulated by 30°/1.0 Hz, a combination that males often perform, as well as 90°/0.5 Hz, which males rarely perform, compared to 10°/0.5 Hz.

## Discussion

Overall, we found that ADLLn responses of female axolotls were strongest when stimulated with a combination of hula parameters that males often displayed during courtship (i.e., 30°/1.0 Hz). However, behavioral responses to hula displays were more broad, such that females exhibited significant responses to more extreme combinations, that males moderately or rarely performed.

### Courtship and Mating Behavior

Hulaing has long been regarded as a male courtship behavior (Halliday, 1990). Nevertheless, we found that female axolotls hula almost as much as males do; importantly, we found that both sexes hulaed for approximately the same total duration during bouts of courtship and mating. Future studies should include female hulaing as an important component of courtship in axolotls and other salamanders.

The characteristics of the male hula changed based on proximity to a female. Males displayed moderate sweep and elevation angles when hulaing in close proximity but used narrow sweep angles and low elevation angles when farther away from the female. Close hula motions likely create more water turbulence, which may serve to elicit a greater response from the lateral line nerves; increased turbulence may also facilitate dispersal of pheromones (Maex et al., 2016; Reddy et al., 2022). In contrast, the parameters that characterize a distant hula may create more laminar flow, which could serve to deliver signals over longer distances.

Females shifted between pausing and locomoting more often when the male was hulaing, which may reflect an increased effort to assess the water disturbances arising from the male’s tail movements. An animal’s own movements generate flow patterns that can interfere with lateral line sensing. In addition, self-motion creates corollary discharge, i.e., motor commands that are routed to sensory systems (Crapse & Sommer, 2016); periods of stillness may mitigate these effects by allowing for unfiltered lateral line input (Lunsford et al., 2019). Perhaps intermittent locomotion is a beneficial strategy for female axolotls trying to localize mates using mechanosensation.

### Behavioral responses to the Robotail

In pilot trials in which male odorants were absent female axolotls routinely attacked the Robotail (**Supplemental Video 1**), which we interpreted as a predatory response. This observation hints at the evolutionary origin of the hula as a type of sensory trap, a phenomenon in which males display courtship signals that mimic other behaviorally relevant stimuli such as prey movements (Christy, 1995). It is also possible that the increased pausing (i.e., locomotion transitions per minute) of females in the presence of fastest and widest tail movements (which may be more prey-like) results from females stopping to discern between prey movements and male cues, particularly as the attacks decreased with the presence of male whole-body odorants. As males rarely perform the combination of wide sweep (90°) with high speed (1.5Hz), these signals along with male odorants may represent a novel cluster of stimuli, and females may pause more to evaluate the unique stimuli. Perhaps male salamanders that exhibit hula-like courtship displays (Houck & Arnold, 2003) have capitalized on an existing behavioral response in females by moving their tails in a way that mimics a prey item.

Given that male axolotls often hulaed at about a 30° sweep angle during live courtship, it is somewhat surprising that, all else being equal, females nudged the base of the tail less often when it was moving at 30° than at 10°. However, because the Robotail could not move freely around the aquarium, it could not provide tactile feedback via “male bumping” as would a live male. During courtship, pairs alternate between male and female bumping, which appears to be important for physically coordinating spermatophore transfer (Shoop, 1960). Perhaps the lack of intermittent and alternating tactile stimulation from an actual male prevented the progression of courtship and reduced non-locomotor courtship behavior.

In analyzing the effects of varying single hula parameters, we found that both sweep angle and speed influenced female locomotion. Females decreased the time spent swimming and walking and decreased the duration of visits to the Robotail in response to wider sweep angles. Faster speeds also resulted in less time swimming but had no effect on walking, and, interestingly, increased the duration of visits to the Robotail and the overall time spent near the Robotail. Wider sweep angles may have prohibited longer visits simply because the Robotail was taking up more space in the designated zone.

In analyzing the effects of combinations of hula parameters, we found that when the Robotail was moving at wider angles and fast speeds, it elicited more frequent transitions between locomoting and pausing. These parameters entail hula movements that are more energetically costly for male axolotls, and in general, more vigorous courtship behaviors are preferred by prospective female mates as signals of mate quality (Byers et al., 2010). Similarly, in our live courtship trials, we found that females transitioned between locomotive states more frequently when the male was hulaing; this suggests that the Robotail may have elicited courtship-like behaviors in females. However, males rarely hulaed using a combination of 90°/1.5 Hz during live courtship trials, which may indicate that females prefer hula motions that are more extreme than what males can typically produce, perhaps an example of receiver bias (Endler & Basolo, 1998, Ryan, 1998) or perhaps a reflection of a general preference for supernormal stimuli or exaggerated features (Bradbury & Vehrenkamp, 2011; Ryan and Cummings, 2013).

### Neurophysiological responses of the ADLLn

In general, we observed larger responses from the ADLLn in female axolotls compared to males. Interestingly, the male ADLLn exhibited both inhibitory and excitatory responses to stimuli, whereas females exhibited only excitatory responses.

Although we only detected marginally significant differences in male ADLLn responses when we varied individual parameters of the Robotail, our small sample size and the relatively low variances among male responses compared to females limits statistical power; therefore, these differences should not be discarded. Interestingly, a sweep angle of 10° and speed of 0.5 Hz elicited inhibitory responses in males but excitatory responses in females. If relatively subtle movements (e.g., 10° sweep angle, 0.5 Hz) were not detected, we would expect the ADLLn to fire at baseline rates; however, in males the ADLLn clearly responded to this stimulus. Interestingly, the difference in male response to 10° and 30° sweep angles was marginally significant and may be biologically meaningful. A similar phenomenon occurs during courtship in *Drosophila melanogaster*; when males detect the presence of male pheromones it triggers an inhibitory response in their P1 neurons, which ultimately prevents occurrences of male-male courtship (Eyjólfsson & Von Philipsborn, 2017). This interplay of inhibitory and excitatory responses and its potential relevance to male-male competition warrants further research.

Females responded more strongly to a sweep angle of 30° than 10°. Given that male axolotls often performed close hula bouts using a moderate (∼30°) sweep angle, this result may indicate a degree of tuning between the female ADLLn and the mechanosensory signals from a nearby courting male. On the other hand, it is somewhat surprising that we detected only marginal differences in female responses to the three speeds we tested, given that behavioral responses differed with hula speed. However, we only recorded activity from the right ADLLn, whereas females in our behavioral assays would have been able to detect mechanosensory stimuli with the right and left ADLLn, plus perhaps other lateral line nerves on the head or even the trunk, in addition to further tuning and modulation that could take place in the central nervous system. Additionally, more vigorous Robotail movements may have created more “noise” as the waves reflected off the walls of the aquarium, and we tested a smaller range of speeds compared to sweep angles; these factors may have reduced the degree of matching between speed and female ADLLn responses.

### Comparison of the male hula to female behavioral and neurophysiological responses

The concordance we observed between female axolotls’ strongest physiological responses and the hula combinations most often performed by males provides support for sender-receiver matching in this system, at least at the level of the nervous system. The greatest excitatory response in the female ADLLn was elicited by the hula combination that males often performed: 30° sweep angle at 1.0 Hz. In contrast, female behavioral responses were somewhat broader. Females exhibited the strongest behavioral responses (i.e., more transitions between locomotor states per min) to three different hula combinations, two of which (30°/1.5 Hz and 90°/1.0 Hz) males performed moderately, and one of which (90°/1.5 Hz) was at the limit of males’ ability and was performed rarely. Thus, our behavioral and neurophysiological results provide support for different models of evolution of communication, indicating that the level of analysis is a critical component in any assessment of such models.

## Supporting information

Supplemental Video 1: Female axolotl attacking the Robotail

## Acknowledgements

Supported by the US National Science Foundation (IOS – 1354089 to HLE), the BEACON Center for the Study of Evolution in Action, and a fellowship from MSU’s Ecology, Evolution, and Behavior Program to TMR. We are extremely grateful to Richard Pease for help with design and fabrication of the Robotail. Thanks to Kay Holekamp and Weiming Li for useful comments on experimental design and data analysis and to Santiago Rodriguez Castro and Samantha Westcott for helpful comments on the manuscript. Special thanks to Laura Muzinic and the Ambystoma Genetic Stock Center at the University of Kentucky (supported by NIH grant P40-OD019794) for providing the animals necessary for these experiments.

## SUPPLEMENTAL MATERIAL

**Supplemental Table 1:**
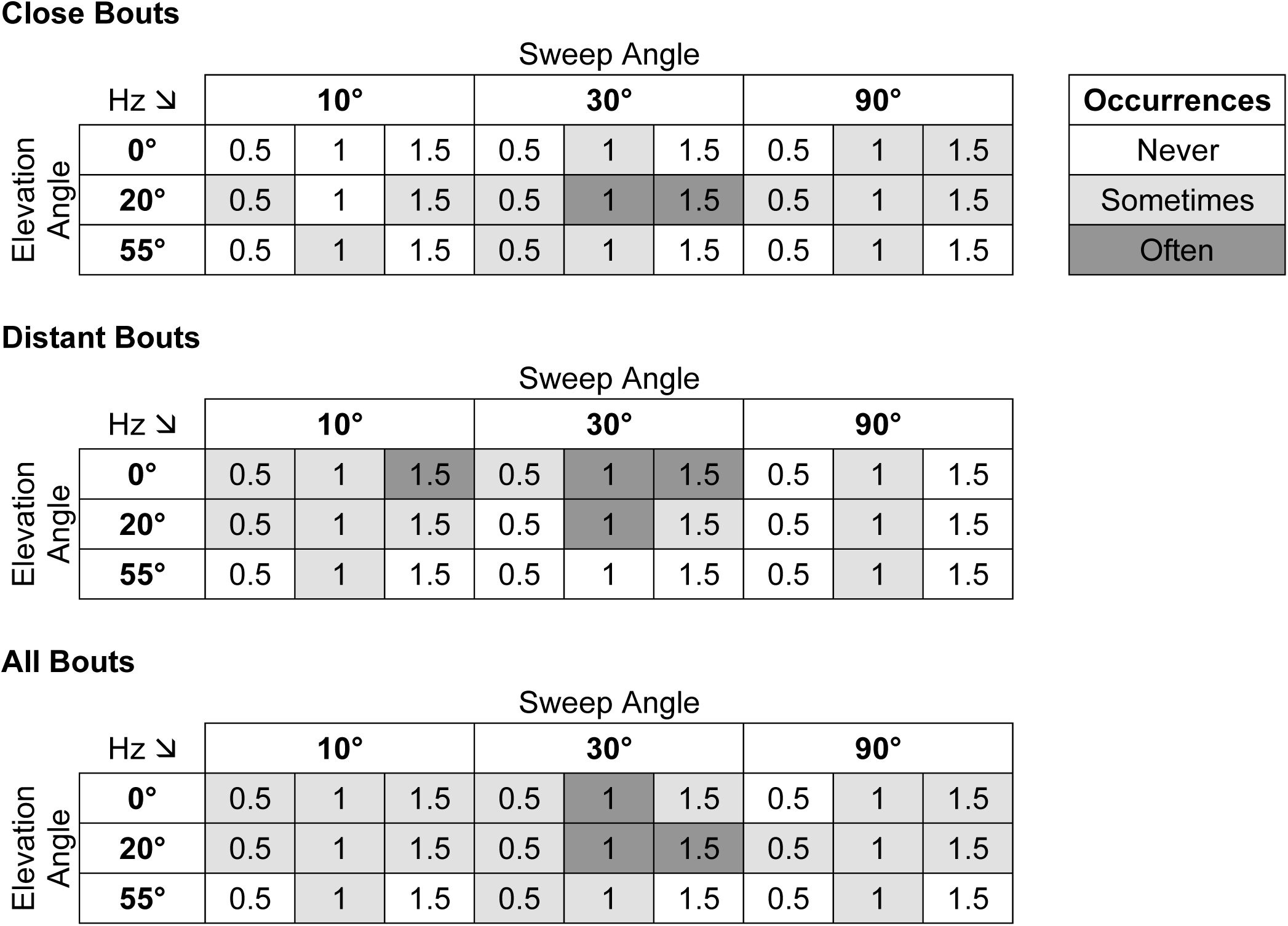
Matrices displaying the occurrence rate of tail motion combinations. Each hula bout was binned and then categorized into one of the 27 motion combinations. Combination frequencies above the median value were deemed as “often”, and values below the median were labeled as “sometimes”, and values of 0 were labeled as “never”.

**Supplemental Table 2:**
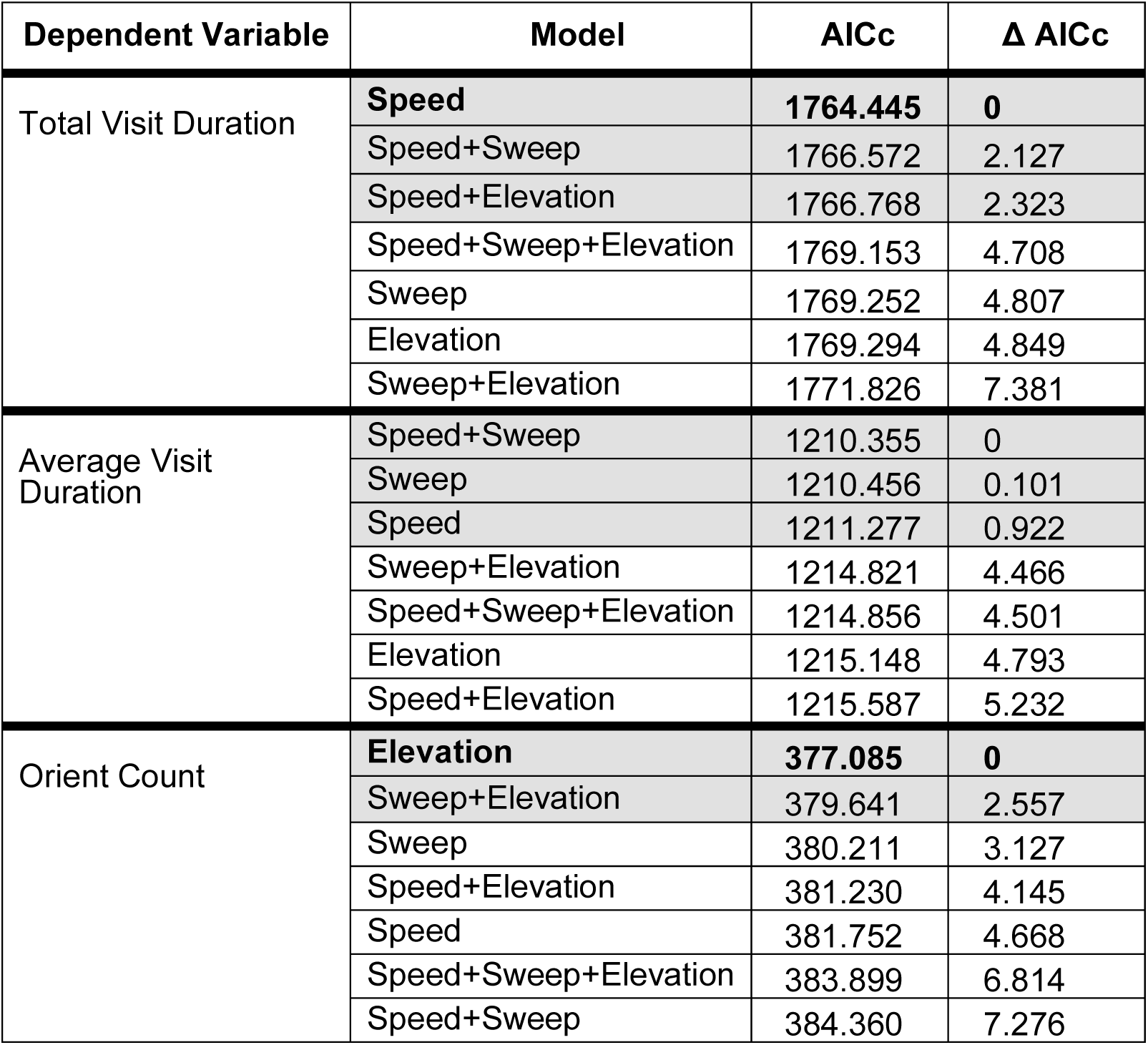
AICc rankings of mixed-effects models for time spent within 1 tail length of the male and orient data. Model selection of seven statistical models to explain total time and average duration per visit of a female to a male as well as the number of times a female orients towards a male. Models in bold are considered the best model based on a delta >3. Models with delta <3 were considered equally weighted and we relied on rules of parsimony to define the best model. Total visit duration was best described by changes in speed, and orient count was best explained by changes in elevation; speed and sweep (alone and in combination) equally explained the average visit duration and there was no clear best model.

**Supplemental Table 3:** AICc rankings of mixed-effects models for physical contacts with the Robotail. Seven statistical models were built for nudge base, nudge tip, and total nudges and their corresponding AICc values calculated to assess which model(s) best explained our data set. Equally weighted models (i.e., within 3 AICc values of the top ranked model) are indicated by gray shading. The model that best described each behavioral variable, which is the simplest model with the lowest AICc value, is indicated by bold text. The model that best explained nudging of the base included just sweep angle, whereas nudging of the tip was best explained by tail elevation; no model best explained the total nudge count.

| Dependent Variable | Model | AICc | $\Delta$ AICc |
| --- | --- | --- | --- |
| Nudge Base Count | **Sweep** | **1119.697** | **0** |
|  | Speed | 1122.730 | 3.033 |
|  | Speed+Sweep | 1123.090 | 3.393 |
|  | Elevation | 1123.168 | 3.471 |
|  | Sweep+Elevation | 1123.644 | 3.947 |
|  | Speed+Elevation | 1126.653 | 6.956 |
|  | Speed+Sweep+Elevation | 1127.155 | 7.458 |
| Nudge Tip Count | Speed+Elevation | 599.138 | 0 |
|  | **Elevation** | **599.704** | **0.566** |
|  | Speed | 602.893 | 3.755 |
|  | Sweep+Elevation | 603.750 | 4.612 |
|  | Speed+Sweep+Elevation | 603.320 | 4.182 |
|  | Speed+Sweep | 606.974 | 7.836 |
|  | Sweep | 607.319 | 8.182 |
| Total Nudge Count | Sweep | 1148.241 | 0 |
|  | Elevation | 1149.645 | 1.404 |
|  | Speed | 1149.646 | 1.405 |
|  | Speed+Sweep | 1151.174 | 2.933 |
|  | Sweep+Elevation | 1151.306 | 3.065 |
|  | Speed+Elevation | 1152.691 | 4.450 |
|  | Speed+Sweep+Elevation | 1154.357 | 6.116 |

**Supplemental Table 4:** Seven statistical models were built for each locomotion variable and their corresponding AICc values calculated to assess which model(s) best explained our data set. Equally weighted models (i.e., within 3 AICc values of the top ranked model) are indicated by gray shading. The model that best described each behavioral variable, which is the simplest model with the lowest AICc value, is indicated by bold text.

| Dependent Variable | Model | AICc | $\Delta$ AICc |
| --- | --- | --- | --- |
| Walk Total Duration | Sweep | 2053.65 | 0 |
|  | Elevation | 2053.98 | 0.337 |
|  | Sweep+Elevation | 2055.66 | 2.018 |
|  | Speed | 2055.76 | 2.112 |
|  | Speed+Sweep | 2057.4 | 3.754 |
|  | Speed+Elevation | 2057.75 | 4.103 |
|  | Speed+Sweep+Elevation | 2059.51 | 5.866 |
| Walk Average Duration | Speed+Sweep | 837.452 | 0 |
|  | **Sweep** | **839.004** | **1.5524** |
|  | Speed+Sweep+Elevation | 839.752 | 2.3002 |
|  | Sweep+Elevation | 841.119 | 3.667 |
|  | Speed | 843.046 | 5.5943 |
|  | Speed+Elevation | 845.565 | 8.1127 |
|  | Elevation | 847.181 | 9.7296 |
| Swim Total Duration | Speed | 1981.95 | 0 |
|  | Sweep | 1982.21 | 0.261 |
|  | Elevation | 1982.52 | 0.567 |
|  | Speed+Sweep | 1985.43 | 3.478 |
|  | Speed+Elevation | 1985.69 | 3.745 |
|  | Sweep+Elevation | 1985.87 | 3.918 |
|  | Speed+Sweep+Elevation | 1989.22 | 7.272 |
| Swim Average Duration | **Sweep** | **1238.13** | **0** |
|  | Speed+Sweep | 1241.04 | 2.906 |
|  | Sweep+Elevation | 1242 | 3.874 |
|  | Speed | 1242.58 | 4.447 |
|  | Elevation | 1244.12 | 5.988 |
|  | Speed+Sweep+Elevation | 1245.05 | 6.921 |
|  | Speed+Elevation | 1246.57 | 8.437 |
| Locomote Total Duration | Speed | 2152.37 | 0 |
|  | Elevation | 2154.48 | 2.115 |
|  | Sweep | 2154.52 | 2.154 |
|  | Speed+Sweep | 2156.16 | 3.795 |
|  | Speed+Elevation | 2156.18 | 3.813 |
|  | Sweep+Elevation | 2158.36 | 5.993 |
|  | Speed+Sweep+Elevation | 2160.13 | 7.758 |
| Locomote Average Duration | Sweep | 1230.63 | 0 |
|  | Speed+Sweep | 1232.01 | 1.379 |
|  | Speed | 1232.53 | 1.9 |
|  | Sweep+Elevation | 1234.14 | 3.506 |
|  | Elevation | 1234.72 | 4.089 |
|  | Speed+Sweep+Elevation | 1235.63 | 5 |
|  | Speed+Elevation | 1236.07 | 5.439 |
| Pause Total Duration | Speed | 2153.56 | 0 |
|  | Sweep | 2155.54 | 1.98 |
|  | Elevation | 2155.56 | 2.003 |
|  | Speed+Sweep | 2157.39 | 3.828 |
|  | Speed+Elevation | 2157.47 | 3.909 |
|  | Sweep+Elevation | 2159.47 | 5.908 |
|  | Speed+Sweep+Elevation | 2161.44 | 7.882 |
| Pause Average Duration | Speed | 1499.97 | 0 |
|  | Sweep | 1501.66 | 1.689 |
|  | Speed+Sweep | 1502.5 | 2.523 |
|  | Elevation | 1502.95 | 2.978 |
|  | Speed+Elevation | 1503.74 | 3.764 |
|  | Sweep+Elevation | 1505.34 | 5.365 |
|  | Speed+Sweep+Elevation | 1506.33 | 6.353 |

**Supplemental Video 1:**
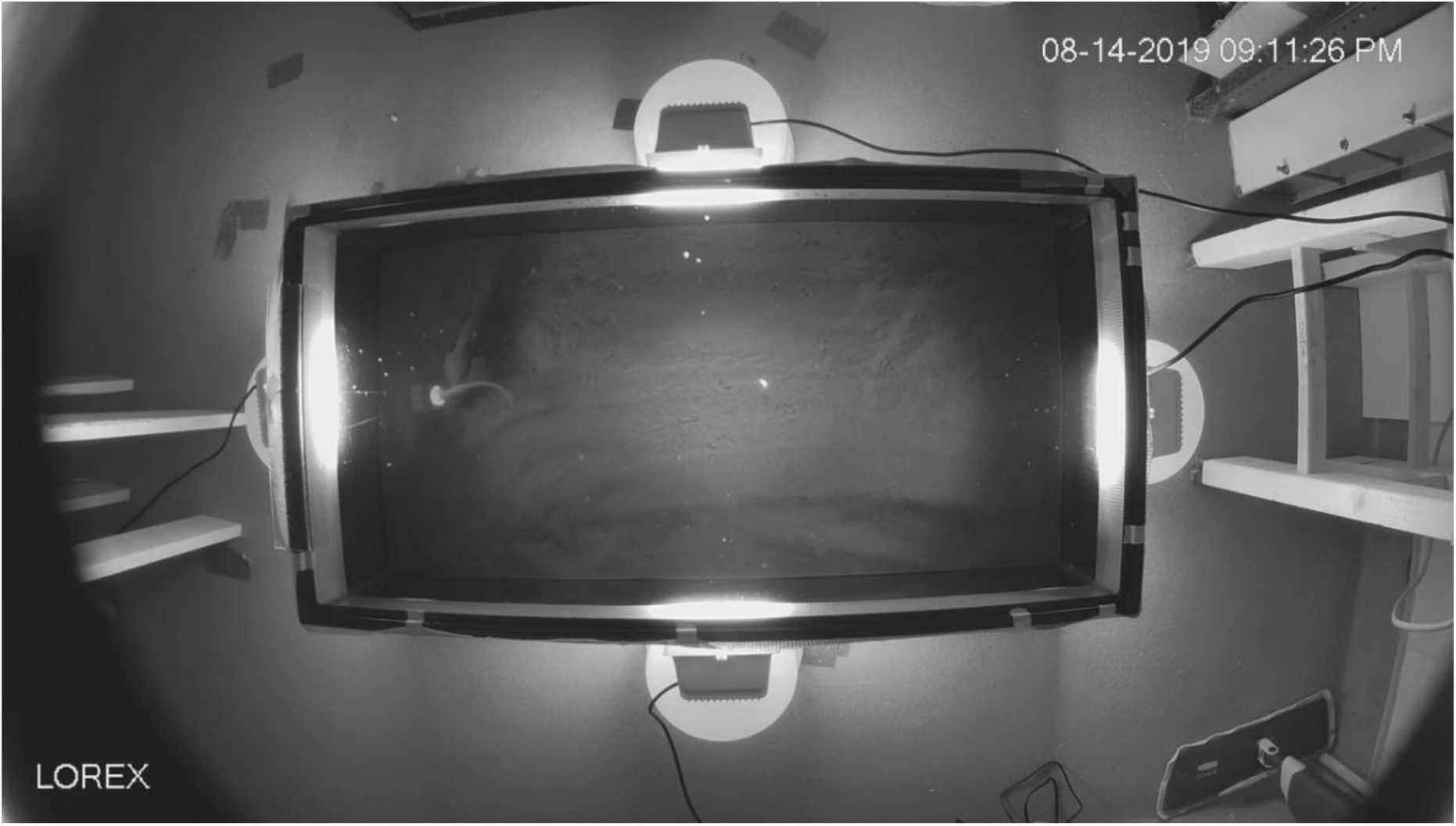
Female axolotl attacking the Robotail. In initial pilot trials, female axolotls attacked the Robotail; we mitigated this problem by introducing male odorants into the test aquarium. In the pilot shown above, the floor of the aquarium is covered in sand; EVA foam tiles were used in subsequent courtship trials. The Robotail is on the viewer’s left and is lighter in color than the adult female axolotl.

